# A standardized method for T cell receptor (TCR) replacement through CRISPR-Cas9 mediated editing and retroviral transduction of primary murine naïve CD8 T cells

**DOI:** 10.64898/2026.08.17.745264

**Authors:** Nadine Tong, John Attanasio, Eric Fagerberg, Kelli Connolly, Nikhil Joshi

## Abstract

CD8 T cells play a central role in immune responses to infection and cancer. However, the diversity of T cell receptor (TCR) specificities makes it challenging to study the mechanisms that regulate T cell activation, differentiation, and effector function. Beyond TCR transgenic mouse models, various complex genome-editing approaches have been employed to overcome this challenge. However, these strategies are often technically demanding, time-intensive, and difficult to adapt. Investigators who are interested in testing *de novo* TCRs under their chosen experimental conditions would benefit from a standardized and accessible method. Here, we describe a protocol that combines ribonucleoprotein (RNP)-based CRISPR-Cas9 editing with retroviral transduction to enable efficient genetic manipulation of murine CD8 T cells. We show that T cells engineered via this protocol can be generated at sufficient scale for downstream *in vitro* assays and *in vivo* adoptive transfer experiments. We expect this method will be useful for investigators who require a standardized and accessible way to study how TCR specificity impacts CD8 T cell responses.

## Introduction

CD8 T cells are known to be central mediators of adaptive immune responses against infection, cancer, and other pathological challenges.^1^ CD8 T cells recognize peptide-major histocompatibility complexes (pMHC) through their T cell receptors (TCRs). Both intrinsic properties of this TCR-pMHC interaction, including TCR specificity and affinity, and extrinsic cues, such as the local cytokine milieu, shape their differentiation into functionally distinct effector and memory populations.^2^ Nevertheless, our understanding of the role that different TCRs play in CD8 T cell differentiation has remained understudied for various reasons, including the difficulty of obtaining sequences for antigen-specific TCRs and the complexity of methods for altering the TCR. The advent and widespread use of single cell RNA sequencing and specifically V(D)J analysis has increased the availability of DNA sequences for antigen-specific TCRs^3^, but the major barrier remains the accessibility of a standardized and reproducible method for generating CD8 T cells expressing these TCRs.

To address this gap, we developed and optimized a rapid and scalable TCR “replacement” protocol that combines CRISPR-Cas9 ribonucleoprotein (RNP)-based editing with retroviral transduction to engineer primary CD8 T cells expressing defined TCRs. Our approach combines and optimizes existing protocols and leverages the ability to use RNP-based editing to efficiently disrupt endogenous TCR expression. The protocol is rapid and reproducible, allowing generation of sufficient cell numbers for both in vitro assays and in vivo adoptive transfer experiments within approximately one week. Moreover, due to the utilization of polyclonal splenocytes from a C57Bl/6 mouse, for most applications this protocol can be easily adapted for different genetic backgrounds, bypassing the need to cross knockout animals to TCR transgenic backgrounds. Finally, because this method relies on standard molecular cloning, retroviral transduction, and CRISPR-Cas9 RNP delivery, it provides a practical framework for replacing TCR specificities in a manner accessible to most laboratories studying CD8 T cell immunology.

### Development of protocol

This protocol combines two established approaches: retroviral transduction of primary T cells^4^ and RNP-based CRISPR-Cas9 editing ^5^ ^6^. By targeting the constant regions of TCRα and TCRβ, we achieve efficient knockout of endogenous TCR chains (>90%) while introducing a full-length TCRαβ pair via retroviral transduction. This design eliminates the need for locus-specific donor templates and bypasses the low efficiency of homology-directed repair (HDR) in primary T cells^7^

Additionally, activation beads are removed before the introduced TCR is expressed, minimizing stimulation bias between cells with high and low levels of transgene expression. We chose Thy1.1 (CD90.1) as a reporter to enable positive selection of successfully transduced cells within four days. The protocol is designed for scalability, accommodating starting cell numbers from 5 x 10^5^ to 4 x 10^6^ per electroporation reaction and producing sufficient engineered T cells for both in vitro and in vivo applications. This method has been used successfully in our lab across multiple TCR specificities and genetic perturbations, enabling rapid testing of multiple TCRs.

### Overview of procedure

This method consists of six stages (Figure 1): (i) isolation of naïve CD8 T cells from murine spleens, (ii) CRISPR-Cas9-RNP electroporation to disrupt endogenous TCR genes, (iii) activation with CD3/CD28 Dynabeads and IL-2, (iv) retroviral transduction with a TCR-encoding construct, (v) magnetic selection of Thy1.1+ cells, and (vi) expansion for downstream assays. The entire procedure requires 6-7 days and yields sufficient cells for in vivo adoptive transfer (1.5-2 × 10 □ CD8 T cells of a desired TCR per 4 × 10□ input cells).

**Figure 1:**
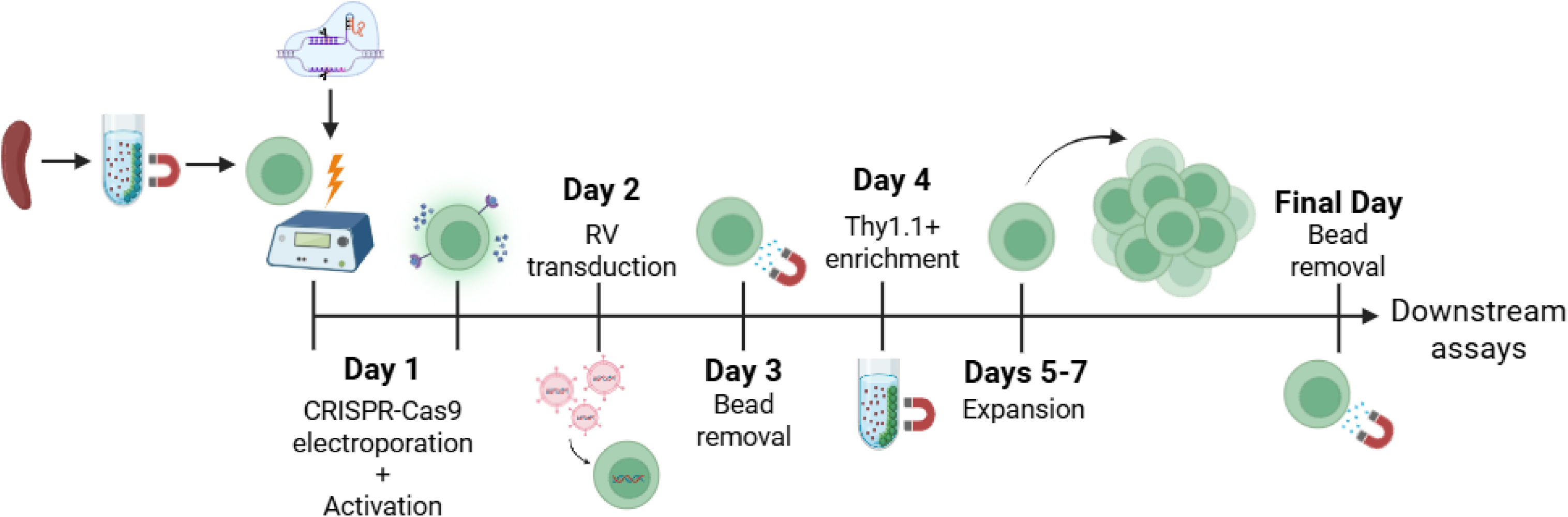
Overview of CRISPR-Cas9-mediated TCR editing and retroviral transduction of primary CD8 T cells. (a) Schematic overview of the workflow. Naïve CD8 T cells are isolated from mouse spleens using negative selection, electroporated with CRISPR-Cas9 RNP complexes targeting the endogenous TRAC and TRBC loci, activated overnight with CD3/CD28 Dynabeads, transduced with a retroviral vector encoding a specific TCR and Thy1.1 reporter, enriched by positive selection, and expanded for downstream applications.

### Applications

#### Studying TCR specificity and antigen recognition

This CRISPR RNP-based approach allows efficient removal of the endogenous TCR while keeping primary T cells healthy and functional. One of its main uses is to test how specific TCRαβ pairs recognize antigen. By knocking out the native TCR chains and reintroducing a defined TCR through retroviral expression, this system isolates pMHC recognition to the engineered TCR alone, minimizing issues from mispaired endogenous chains. This makes it possible to attribute functional readouts to the introduced receptor.

#### TCR signaling and effector differentiation studies

Because the edited CD8 T cells remain viable and expand well, they can be used to study how antigen stimulation shapes downstream signaling and differentiation. The Thy 1.1 reporter is useful here, since it allows clear identification and enrichment of engineered cells.

#### In vivo adoptive transfer into infection and tumor models

The number of cells generated with this method (1.5–2 × 10 cells per 4 × 10 input cells) is sufficient for adoptive transfer into standard mouse models. These engineered T cells can be transferred into congenic hosts and tracked over time in blood, lymphoid organs, and tissues using anti-Thy1.1 antibodies, consistent with established adoptive transfer approaches^8^. Because they remain fully functional in vivo, they can be used to study antigen-driven expansion, trafficking, and effector responses in both infection and tumor settings, enabling controlled comparisons across different TCRs and antigens.

#### Screening multiple TCRs in parallel

The short timeline (6-7 days) and lack of complex donor template design make this system well-suited for parallel screening of multiple TCRs. Different retroviral constructs can be introduced into separate cell populations after a shared CRISPR knockout step, allowing multiple TCRs to be evaluated side-by-side under the same experimental conditions for direct functional comparisons.

### Comparison to other methods

As mentioned above, several alternative strategies exist for studying defined TCRs in murine CD8 T cells. Retrogenic approaches have allowed for the generation of T cells expressing defined TCRs without the need to create new transgenic lines by retroviral transduction of hematopoietic stem and progenitor cells isolated from donor bone marrow ^9^. This system enables the study of defined TCRs without the need for germline engineering and allows for relatively flexible manipulation of gene expression ^10^. However, the requirement for irradiation, bone marrow reconstitution, and in vivo development introduces significant experimental complexity and extended timelines, often requiring several weeks to months before usable T cells can be obtained ^9^.

Orthotopic TCR replacement strategies extend this concept by introducing transgenic TCR sequences into endogenous loci, such as the TRAC locus, placing the introduced receptor under native transcriptional regulation ^11^. This strategy minimizes mispairing between endogenous and introduced TCR chains and produces more physiologically regulated receptor expression. However, these strategies depend on HDR, requiring efficient donor template delivery and careful optimization of editing conditions in primary T cells. Recent advances have begun to address these limitations through non-viral delivery platforms and improved selection strategies ^12^, using small molecules that bias DNA repair towards HDR, including SCR7 and RS-1^13^. Despite these improvements, orthotopic replacement strategies remain technically challenging and are not yet easily scalable across many laboratories.

More recently, adeno-associated virus (AAV)-mediated genome editing approaches have provided a more efficient and targeted strategy for genetic manipulation of primary T cells ^14,15^. This strategy supports targeted integration, multiplexed editing, and delivery of large DNA sequences ^16^, and has been applied in both experimental and therapeutic settings. Despite this, AAV-based strategies also rely on HDR, requiring co-delivery of donor templates matched to the target locus, which increases design and delivery complexity ^15^. In addition, AAV-based approaches require specialized viral production infrastructure ^17^ and are subject to extensive institutional biosafety and regulatory oversight, often requiring approval from Institutional Biosafety Committees (IBCs) and adherence to NIH and FDA gene therapy guidelines. This further complicates and delays experimental implementation and may limit accessibility for some laboratories.

In contrast to the systems described above, our TCR “replacement” protocol provides a rapid workflow for engineering primary CD8 T cells expressing defined TCRs, without the need for complex donor template design or time-intensive optimization of HDR-based editing, and instead relies on techniques already widely used in immunology laboratories. It enables efficient removal of endogenous TCRs while introducing transgenic receptors in a single coordinated procedure completed in approximately one week. This design supports flexible testing of diverse TCR specificities under uniform experimental conditions, while maintaining robust cell recovery and viability. While not intended to supplant the methods described above, the method is intended to be straightforward and easy to implement. Depending on the specific needs of the question being asked, we envision the TCR replacement method will be useful for many investigators as a means towards asking questions, screening TCRs of interest, and/or narrowing pools of TCRs for downstream applications, such as retrogenic animals or AAV-based vector generation.

### Experimental design

#### Identifying TCR sequences

Our method relies on the identification and cloning of a TCRαβ pair of interest, which can be obtained through a variety of methods. This is an active area of research and other publications have described methods for the identification of antigen specific TCRs and recovery of paired TCRαβ sequences ^18^ ^19^ ^20^. For this demonstration, we use the P14 TCR that is specific for the lymphocytic choriomeningitis virus (LCMV) glycoprotein (GP) peptide 33-43 (GP33)^21^.

#### Cloning/ Vector Preparation

The TCR of interest is subcloned into the pMIG-II vector (Addgene, cat.no. 52107) using standard restriction enzyme-based and Gibson assembly methods. The final construct replaces the native IRES-eGFP cassette with Thy1.1 (CD90.1), a cell-surface reporter that remains detectable following fixation and permeabilization procedures commonly used for intracellular staining of T cell differentiation markers. The pMIG-II backbone is linearized using EcoRI-HF (New England Biolabs, cat. no. R3101S) and NotI-HF (NEB, cat. no. R3189S) restriction enzymes in rCutSmart buffer (NEB, cat. no. B6004S), after which the TCR insert and Thy 1.1 reporter fragments are assembled into the vector via Gibson assembly (NEB, cat. no. E2611S). Silent mutations are introduced into PAM recognition sequences within the cloned TCR coding regions as a safeguard against Cas9-mediated deletion after editing. Following assembly, constructs are verified by gel electrophoresis, subject to bacterial transformation, and purified by miniprep (Qiagen, cat. no. 27106) prior to downstream applications and retroviral production.

Purified plasmid constructs are stored at 4°C for short-term use.

#### gRNA design, preparation, and storage

We target the constant regions of TCRα (TRAC) and TCRb (TRBC) using validated sgRNA sequences:

TRAC: CGUCAUGAGCAGGUUAAAUC

TRBC: CCCACCAGCUCAGCUCCACG

sgRNAs are custom ordered (e.g., from Synthego). Guides should be spun down at >10,000g for 5 minutes and resuspended at 300 µM in nuclease-free water. We strongly encourage aliquoting guides as single-use reactions to avoid freeze-thaw degradation. Aliquots are stored at –80 °C.

#### Retroviral production

Retroviral supernatant is produced by transfecting Plat E cells. Plat E cells are cultured to 70% confluency in Plat E media. Cells are transfected using Lipofectamine 2000 according to the manufacturer’s instructions. Viral supernatant is harvested 48-72 hours post-transfection, pooled, aliquoted, and stored at –80 °C until use. We strongly encourage users to optimize this step for themselves, as different vectors and methods of producing virus will yield different results. For consistent results, we recommend producing a single large batch of supernatant for a given TCR and using it across all experiments to minimize batch-to-batch variability.

#### Cell number and scaling

The protocol is optimized for 4 × 10 CD8 T cells per electroporation well, yielding approximately 1.5–2 × 10 Thy1.1+ cells after 7 days. Successful scaling down to 5 × 10 cells is possible but reduces absolute yield. Scaling below 1 × 10□ cells per well is not recommended because cell loss during magnetic enrichment steps becomes limiting. When scaling up, we advise performing multiple parallel electroporation reactions rather than using larger cassettes (e.g., Lonza 100-µL strips), as we have observed reduced viability with larger reaction volumes. No more than two spleens (approximately 12-20 × 10 total CD8 T cells) should be combined per isolation reaction to maintain purity (>95% CD8).

#### Stimulation and transduction parameters

Activation with CD3/CD28 Dynabeads is essential for both retroviral transduction and efficient RNP editing. We use a 1:1 bead-to-cell ratio; lower ratios may be used for specific expansion goals, but could impair transduction efficiency, while significantly higher ratios cause overstimulation and exhaustion. IL-2 supplementation at 5 ng/mL is optimal. Transduction is performed 20-22 hours after activation, which corresponds to peak CD8 T cell activation. Strong clustering around Dynabeads should be observed. Spinfection at 600g for 1.5 h at 32 °C with protamine sulfate (8 µg/mL) consistently yields higher efficiency than static transduction.

#### Controls and validation

To validate the functionality and physiological relevance of our method, T cells generated using this protocol were directly compared to CD8 T cells isolated from transgenic P14 mice, which express a well-characterized TCR specific for the LCMV GP33 epitope. P14 T cells served as a positive control for antigen-specific TCR expression, activation, and effector differentiation.

Engineered T cells were assessed alongside P14 controls for TCR expression, transduction efficiency, differentiation, and antigen-specific responses.

### Limitations

Several limitations of this protocol should be considered.

First, retroviral transduction efficiency is dependent on viral titer, which can vary substantially between viral preparations due to differences in packaging efficiency, plasmid quality, or handling conditions. As a result, inconsistent transduction rates may introduce experimental variability in transgene expression levels and reduce reproducibility across experiments. To mitigate this concern, we have utilized a construct containing a co-translationally expressed Thy 1.1 reporter downstream of a 2A sequence, which allows us to compare transduced T cells with equal retroviral expression (*i.e.*, between populations transduced with different constructs).

In addition, although CRISPR-mediated knockout efficiency is typically high, incomplete editing can still occur in some cells, leading to residual endogenous TCR α or β chain expression. This residual expression increases the possibility of mixed TCR pairing between endogenous and transgenic chains, altering receptor specificity and uniformity, and/or complicating the interpretation of downstream functional assays.

Another limitation is that the specific TCR is introduced through retroviral transduction rather than targeted insertion into the endogenous TCR locus. Consequently, TCR expression is driven by exogenous regulatory elements and does not fully recapitulate endogenous transcriptional control or physiological surface expression dynamics of the TCR locus.

Finally, although the present protocol is optimized for rapid expansion and high cellular yield using IL-2, incorporation of additional cytokines, such as IL7 and IL15, may further improve the phenotypic and functional properties of the resulting T cell product depending on the intended downstream application.

### Regulatory approvals

This protocol involves the use of live mice (C57BL/6, 6–10 weeks old). All procedures must be approved by the Institutional Animal Care and Use Committee (IACUC) or equivalent ethical review body.

## Materials

### REAGENTS

- Cell culture:

- RPMI-1640 with L-Glutamine (Gibco, cat.no.11875-093)
- DMEM, 1x, + 4.5 g/L D-Glucose, + L-Glutamine, + 110 mg/L Sodium Pyruvate (Gibco, cat.no. 11995065)
- DPBS, 1×, no Calcium Chloride, no Magnesium Chloride (Gibco, cat.no. 14190-144)
- EDTA, 0.5 M Solution pH 8.0 (RPI, cat.no. E14000-500.0)
- Fetal Bovine Serum (FBS) (Sigma-Aldrich, cat.no. F0926)
- Penicillin–streptomycin (Gibco, cat.no. 15140-122)
- GlutaMAX™ Supplement 100X (Gibco, cat.no. 35050-061)
- Antibiotic-Antimycotic 100X (Gibco, cat.no. 15240-062)
- Hepes 1M (Gibco, cat.no. 15630-080)
- 2-Mercaptoethanol (Sigma Aldrich, cat.no. M3148) **<u>CAUTION:</u>** 2-Mercaptoethanol is toxic upon inhalation or contact with skin and is harmful if swallowed. Handle with proper protective equipment under a fume hood.
- Nuclease-free Water (Synthego, cat. no. #0000000049)
- 1X RBC Lysis Buffer (eBioscience, cat.no. 00-4333-57)
- EasySep Mouse CD8+ T Cell Isolation Kit (StemCell, cat.no.19853)
- EasySep Mouse CD90.1+ T Cell Isolation Kit (StemCell, cat.no.18958)
- P3 Primary Cell 4D-Nucleofector® X Kit S (Lonza, cat.no. V4XP-3032)
- Human IL-2 Protein, PeproTech® (Gibco, cat.no. 200-02-50UG)
- Protamine sulfate (Sigma Aldrich, cat. no. P3369-10G)
- Trypan Blue Stain 0.4% (Gibco, cat.no. 15250-061)
- Dynabeads Mouse T-Activator CD3/CD28 (Gibco, cat.no. 11452D)
- Cas9 Nuclease V3, 500 µg (IDT, cat.no. 1081059)
- Custom sgRNA (Synthego)
  - TRAC CGUCAUGAGCAGGUUAAAUC
  - TRBC CCCACCAGCUCAGCUCCACG
- Plastics:

- 5 mL polystyrene round-bottom tube (BD Falcon, cat.no. 352052)
- 50 mL polypropylene conical tube (BD Falcon, cat.no. 352070)
- 15 mL polypropylene conical tube (BD Falcon, cat.no. 352096)
- 1 mL TB Syringe (BD, cat.no. 309659)
- 3 mL TB Syringe (BD, cat.no. 309657)
- 12-well plate, tissue-culture-treated (Corning, cat.no. 3513)
- 6-well plate, tissue-culture-treated (BD Falcon, cat.no. 353046)
- 25cm^3^ Polystyrene Tissue Culture Flask, Vented Cap (Falcon, cat. no. 353109)
- 70-μm cell strainer (BD Falcon, cat.no. 352350)
- 0.2 mL polypropylene PCR 8-Tube Strips (USA Scientific, cat.no. 1402-4700)
- 5 mL polypropylene round-bottom tube with cell strainer cap (BD Falcon, cat.no. 352235)
- 10 mL, 5 mL serological sterile pipettes (CELLTREAT, cat.no. 229210, 229205)

### EQUIPMENT

- EasyStep Magnet (StemCell, cat.no. 18000)
- Vortex Mixer (Scientific Industries, cat.no. SI-0236)
- Centrifuge (Eppendorf, cat.no. 5948FI001388)
- 4D-Nucleofector™ Core Unit (Lonza, cat.no. AAF-1001B)
- 4D-Nucleofector™ X Unit (Lonza, cat.no. AAF-1001X)
- Flow cytometer analyzer (BD Biosciences, model no. BD FACSymphony)

### REAGENT SETUP

- Plat E Media (Store at 4 °C)

- mL DMEM 1x
- 10% FBS
- 1% Anti-Anti
- 1% Pen Strep
- T Cell Media (Store at 4 °C)

- mL RPMI-1640
- 10% FBS
- 1% Anti-Anti
- 1% Pen Strep
- 1% GlutaMAX
- mL of Hepes
- µL of 2-Mercaptoethanol
- MACS Buffer (Store at 4 °C)

- mL DPBS 1X
- 0.5-2% FBS
- µL EDTA

#### Preparation

Cloning - The T cell receptor of interest was subcloned into the pMIG II retroviral vector, in which IRES-eGFP was replaced with 2A-Thy1.1 to enable improved reporter retention following fixation and permeabilization. Silent mutations were introduced into the PAM recognition sequences of the cloned TCR as a safeguard against Cas9 mediated deletion.

Retroviral Production - Plat E cells were cultured to 70% confluency and transfected using Lipofectamine 2000 according to the manufacturer’s instructions. Viral supernatant was harvested 48-72h post-transduction, aliquoted, and stored at −80°C until use. We strongly encourage users to optimize this step for themselves as different vectors and methods of producing virus will yield different results.

**CAUTION:** Standard precautions must be taken while handling retroviral supernatants, and all experiments should be carried out in at least class II biological safety cabinets (or as required by institutional guidelines) and using appropriate protective equipment.

sgRNAs – sgRNAs were custom ordered through Synthego. When receiving guides, they should be spun down at >10000g for 5 minutes and resuspended at 300 µM. We strongly encourage aliquoting guides as single use reactions to avoid waste.

**Note:** This protocol was optimized for use with a 16-well Lonza nucleofection cassette and a maximum of 4 x 10^6^ cells per well.

#### Day 1: Mouse Harvest (20 minutes)

**<u>CRITICAL:</u>** It is critical to have a pure population (>95%) of CD8 T cells for subsequent days. An 8-week-old C57BL/6 mouse yields approximately 6 x 10^6^ – 10 x 10^6^ CD8 T cells. If higher cell numbers are required, no more than two spleens should be combined per isolation reaction.

1. Euthanize mice in accordance with institutional animal care and use guidelines.
2. Spray 70% ethanol over the mouse body, and collect spleens in appropriately sized tubes submerged in MACS buffer on ice.
3. Pour off spleen and MACS buffer onto a 70 µm sterile cell strainer placed over a 50 mL conical tube.
4. Crush spleen through strainer using the plunger of a 1 mL or 3 mL syringe.
5. Wash the strainer twice with 10 mL of MACS buffer.
6. Centrifuge the 50 mL tube at 450g for 5 minutes at 4°C.
7. Aspirate the supernatant and resuspend the cell pellet in 2 mL of RBC lysis buffer.
8. Incubate at room temperature for 3-5 minutes.
9. Quench lysis by adding 10X volume of MACS buffer.
10. Centrifuge the 50 mL tube at 450g for 5 minutes at 4°C.
11. Resuspend the cell pellet in 1 mL of MACS Buffer.

#### Day 1: Cell Isolation (20 minutes)

12. Enrich CD8 T cells by CD8 negative selection according to the manufacturer’s protocol.
13. Pour off negatively selected CD8 T cells into a fresh 15 mL conical tube.
14. Bring volume of isolated CD8 cell suspension to 15 mL with MACS buffer.
15. Centrifuge the 15 mL tube at 450g for 5 minutes at 4°C.
16. Aspirate the supernatant.
17. Resuspend cells at approximately 2 x 10^6^ cells/mL for counting. **Note:** Resuspending the cell pellet in 5 mL of buffer per spleen yields an optimal concentration for counting.
18. Count using preferred cell counting method.
19. Place cells on ice.
20. Proceed to step 21.

#### Day 1: Assembly of Ribonucleoprotein (RNP) complexes (30 minutes)

**<u>CRITICAL:</u>** Guides targeting the TCRα and TCRβ constant regions must be complexed separately. Premature mixing of sgRNAs may reduce efficient RNP formation. They will later be combined into a single reaction.

21. Per reaction, combine 0.5 µL of Cas9 nuclease (5µg), 1 µL of nuclease-free water, and 1 µL of sgRNA into a PCR tube.
22. Incubate at room temperature for 20-45 minutes.

#### Day 1: Cell Electroporation (20 minutes)

**<u>Reminder:</u>** This protocol was optimized for use with a 16-well Lonza nucleofection cassette and a maximum of 4 x 10^6^ cells per well. We have electroporated as low as 5 x 10^5^ cells.

23. Take desired cell number from step 19 and transfer into a new 15 mL tube.
24. Top up the 15 mL conical with isolated CD8 T cells with 1x DPBS.
25. Centrifuge at 450 g for 5 minutes at 4°C.
26. During the first wash, prepare electroporation buffer by adding 16 µL of the Lonza P3 buffer and 4.5 µL P1 supplement to an Eppendorf tube.

**<u>CRITICAL:</u>** If performing multiple reactions, maintain this P3/P1 ratio. Altering this ratio may reduce electroporation efficiency and cell viability.

27. Set up the Lonza 4D-Nucleofector instrument.
28. Select the drive (The 16-well cassette should be pictured).
29. Select wells to be used on the screen.
30. Choose cell type program (“T cell, mouse, unstim.” If not preset, manually enter DN100).
31. Select Start/Open. Leave open and return to cells.
32. Aspirate the supernatant and repeat DPBS 1x wash (Step 23-24).
33. During the second wash, combine RNP complexes with electroporation buffer for total volume of 25.5 µL.
34. Return to cells and aspirate the supernatant completely.

**<u>CRITICAL:</u>** It is critical to remove all supernatant for optimal electroporation efficiency. Use a pipette to remove any liquid if necessary.

35. Resuspend cells in 25 µL of the RNP-containing P3/P1 buffer per reaction.

**<u>CRITICAL:</u>** Resuspend slowly to avoid bubble formation. Cell viability rapidly decreases in P3/P1 buffer; subsequent steps should be performed quickly.

36. Transfer cell suspension to the Lonza 16-well cassette.
37. Place cassette in the open tray of the instrument and click start.
38. Remove cassette immediately from instrument tray upon program completion.
39. Add 125 µL of pre-warmed T cell media to each well.
40. Incubate at 37°C for 10 minutes.
41. Transfer cells to a FACS tube, top up with additional pre-warmed T cell media.
42. Centrifuge at 450g for 5 minutes at 4°C.
43. Resuspend cells at approximately 2 x 10^6^ cells/mL in T cell media for counting. **Note:** 25-50% cell loss during electroporation is normal.
44. Transfer cells into a tissue culture plate. Use the table below as a guide for seeding densities of CD8 T cells during stimulation. 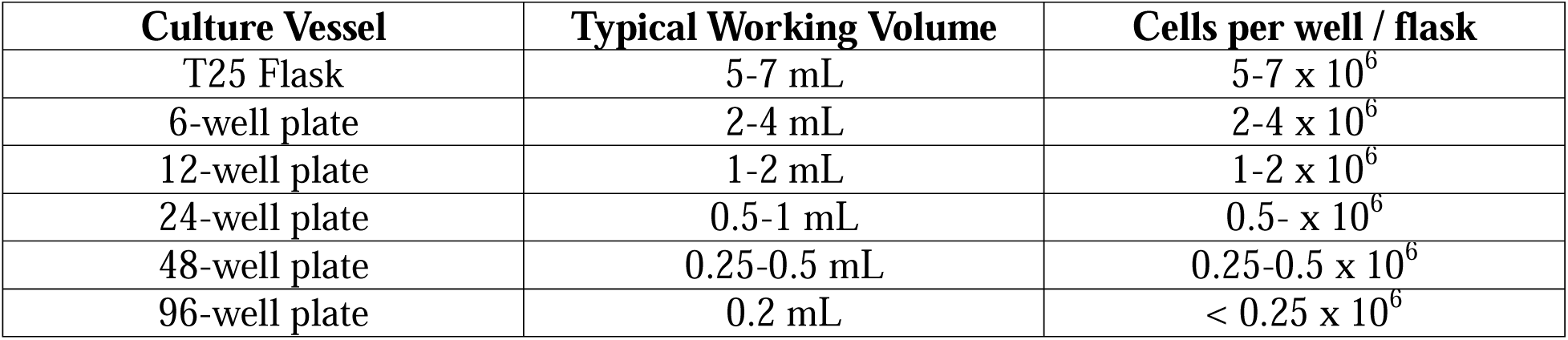
45. Stimulate with CD3/CD28 Dynabeads at a 1:1 bead-to-cell ratio and supplement with IL-2 (5ng/mL final dilution).
46. Incubate at 37°C overnight.

#### Day 2: Retroviral Transduction (2 hours)

**CRITICAL:** Optimal CD8 T cell activation with Dynabeads occurs after 20-22 hours.

**Note:** We perform transductions using standardized volumes of viral supernatant generated in large batches to minimize variability between experiments.

**<u>CAUTION:</u>** All retroviral work should be performed in at least a class II biosafety cabinet with appropriate personal protective equipment. Retrovirus must be handled and disposed of in accordance with your institutional biohazard regulations.

47. Pre-warm centrifuge to 32°C.
48. Harvest cells into a FACS tube.
49. Centrifuge at 450g for 5 minutes at 4°C.
50. Aspirate the supernatant.
51. Resuspend the cell pellet in the appropriate volume of viral supernatant. Use the table below as a guide. 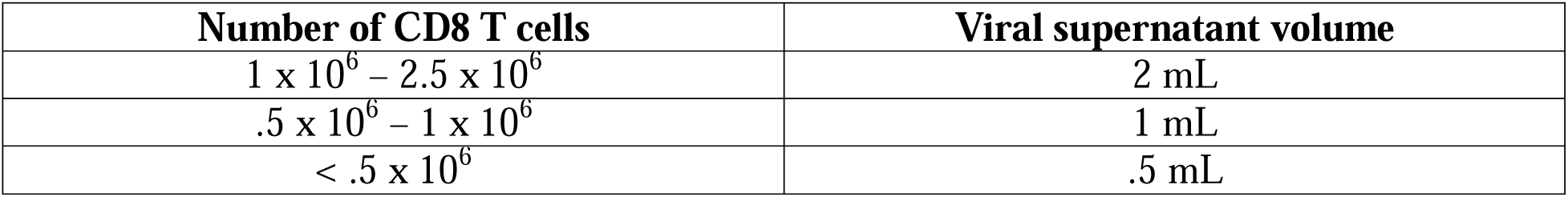
52. Supplement with IL-2 (5ng/mL final dilution) and protamine sulfate (8µg/mL final dilution).
53. Resuspend cells and transfer to an appropriately sized tissue culture plate.
54. Spinfect (centrifugation-enhanced viral transduction) at 600g for 1.5 hours at 32°C with low acceleration/deceleration and bucket lids secured.
55. Transfer plate to a 37°C incubator and incubate overnight.

#### Day 3: Dynabead Removal (15 minutes)

**CRITICAL:** Continued CD3/CD28 stimulation beyond this point is unnecessary for transduction and may promote excessive activation and differentiation, reducing cell viability and increasing experimental variability.

56. Harvest cells into a FACS tube.
57. Centrifuge at 450g for 5 minutes at 4°C.
58. Aspirate supernatant and resuspend in 2.5 mL of MACS Buffer.
59. Place tube into the EasySep Magnet for 3 minutes at room temperature.
60. Pour off the supernatant containing bead-free cells into a new FACS tube.
61. Centrifuge at 450g for 5 minutes at 4°C.
62. Aspirate supernatant.
63. Resuspend in prewarmed T cell media with IL-2 (5ng/mL final dilution).
64. Transfer cells to an appropriately sized tissue culture plate.
65. Incubate overnight at 37°C.

#### Day 4: Thy 1.1+ Selection (30 minutes)

**CRITICAL:** Early enrichment removes untransduced cells before prolonged culture, reducing competition for cytokines and generating a purer transduced population for downstream expansion.

66. Harvest cells into a FACS tube.
67. Centrifuge at 450g for 5 minutes at 4°C.
68. Aspirate supernatant and resuspend in 2.5 mL of MACS Buffer.
69. Enrich Thy1.1+ cells by Thy1.1 positive selection per the manufacturer’s protocols.
70. After selection, pour off unwanted cells.
71. Top up FACS tube with MACS buffer.
72. Centrifuge at 450g for 5 minutes at 4°C.
73. Aspirate the supernatant and resuspend in pre-warmed T cell media.
74. Count.

**<u>Note:</u>** Under our standard viral production conditions, transduction efficiencies consistently range from 20–30%. Alternative viral production methods or viral concentration may change the yield of selected cells.

75. Plate cells in an appropriately sized tissue culture plate at 1 x 10^6^ cells/mL of T cell media supplemented with IL-2 (5ng/mL final dilution). **Note:** We typically downsize by two plate sizes following selection to maintain optimal cell density.
76. Incubate overnight at 37°C.

#### Days 5-7: Cell Maintenance (10 minutes)

77. Harvest cells into a FACS tube.
78. Centrifuge at 450g for 5 minutes at 4°C.
79. Aspirate supernatant and resuspend pellet in pre-warmed T cell media.
80. Count cells.
81. Replate cells in an appropriately sized tissue culture plate at 1 x 10^6^ cells/mL of T cell media supplemented with IL-2 (5ng/mL final dilution).
82. Incubate overnight at 37°C.

**Note:** Repeat daily as needed to maintain optimal cell density and viability.

#### Final Day: Removal of Thy1.1 Selection Beads (10 minutes)

83. Harvest cells into a FACS tube.
84. Centrifuge at 450g for 5 minutes at 4°C.
85. Aspirate the supernatant and resuspend in 2.5 mL of MACS Buffer.
86. Place tube into the EasySep Magnet for 3 minutes at room temperature.
87. Pour off the supernatant containing bead-free cells into a fresh FACS tube.
88. Centrifuge at 450g for 5 minutes at 4°C.
89. Aspirate supernatant and resuspend cells in pre-warmed T cell media.
90. Count cells. **Note:** If ∼4 x 10^6^ cells were used for electroporation and subsequent steps, the expected final yield is approximately 1.5 x 10^7^ – 2 x 10^7^ Thy 1.1^+^ T cells.
91. Cells are now ready for downstream assays.

#### Timing

- Day 1: 3-4 hours

- Steps 1-11, 20 minutes
- Steps 12-20, 20 minutes
- Steps 21-22, 30 minutes
- Steps 23-46, 1.5-2 hours
- Day 2:

- **S**teps 47-55: 2 hours
- Day 3:

- Steps 56-65: 15-20 minutes
- Day 4:

- Steps 66-76: 30-45 minutes
- Days 5 – 7:

- Steps 77-82: 10 – 15 minutes per day
- Final Day:

- Steps 83-91: 10 – 15 minutes

#### Troubleshooting

Troubleshooting:
| Steps | Problem | Possible Reason | Solution |
| --- | --- | --- | --- |
| 37 | Low CRISPR editing efficiency | Improper RNP assembly; degraded sgRNA; incorrect nucleofection program | Use freshly thawed sgRNA aliquots and avoid repeated freeze-thaw cycles.<br>Confirm proper RNP incubation time before electroporation.<br>Verify nucleofection program and that reagents are prepared at the recommended concentrations. |
| 45-46 | Poor activation or expansion following | Incorrect bead-to-cell ratio; insufficient IL-2 supplementation; low | Use a 1:1 Dynabead-to-cell ratio and confirm IL-2 is added at 5ng/mL final concentration. |
|  | stimulation. | starting cell viability | Adding up to 10 ng/mL IL-2 could also improve cell expansion, but avoid overstimulation. |
| 51-55 | Low retroviral transduction efficiency | Low viral titer; suboptimal activation timing | Use freshly thawed high-titer viral supernatant whenever possible.<br>Perform transduction 20-22 hrs after activation. |
| 90 | Final cell yield lower than expected | Cell loss during electroporation or enrichment; poor viral transduction; cell death during culture | Confirm viability after electroporation and monitor culture daily.<br>Follow previous troubleshooting recommendations. |

#### Anticipated Results

Upon completing the procedure, cells should express the introduced transgenic TCR and the Thy 1.1 reporter. In our example, the retroviral construct encodes the P14 TCRα and TCRβ chains, separated by 2A sequences, and with a co-translationally expressed 2A-Thy1.1 (note that the nucleotide sequences of two 2A sequences are degenerate relative to each other). Successful TCR replacement can then by assessed with FACS analyses by staining cells for Thy1.1 (clone OX-7), CD8α (clone 53-6.7), GP33 Tetramer (H2-Db LCMV GP var C41M 33-41; NIH Tetramer core facility), pan-TCRβ (clone H57-597) and Vα2 (clone B20.1). It is important to note that the representative data shown in the following figures were obtained from multiple independent experiments performed to validate different stages of the protocol.

The first critical checkpoint is efficient CRISPR-Cas9-mediated disruption of the endogenous TCRα and TCRβ chains. In figure 2, OT-1 cells were used to demonstrate effective disruption of endogenous expressing TCR chains. Electroporation of TRAC- and TRBC-targeting sgRNAs should result in near-complete loss of Vα2 and TCRβ surface expression compared to cells electroporated without guide RNAs. Because efficient endogenous TCR disruption is essential for successful receptor replacement, incomplete knockout represents a major point of failure for the protocol.

**Figure 2:**
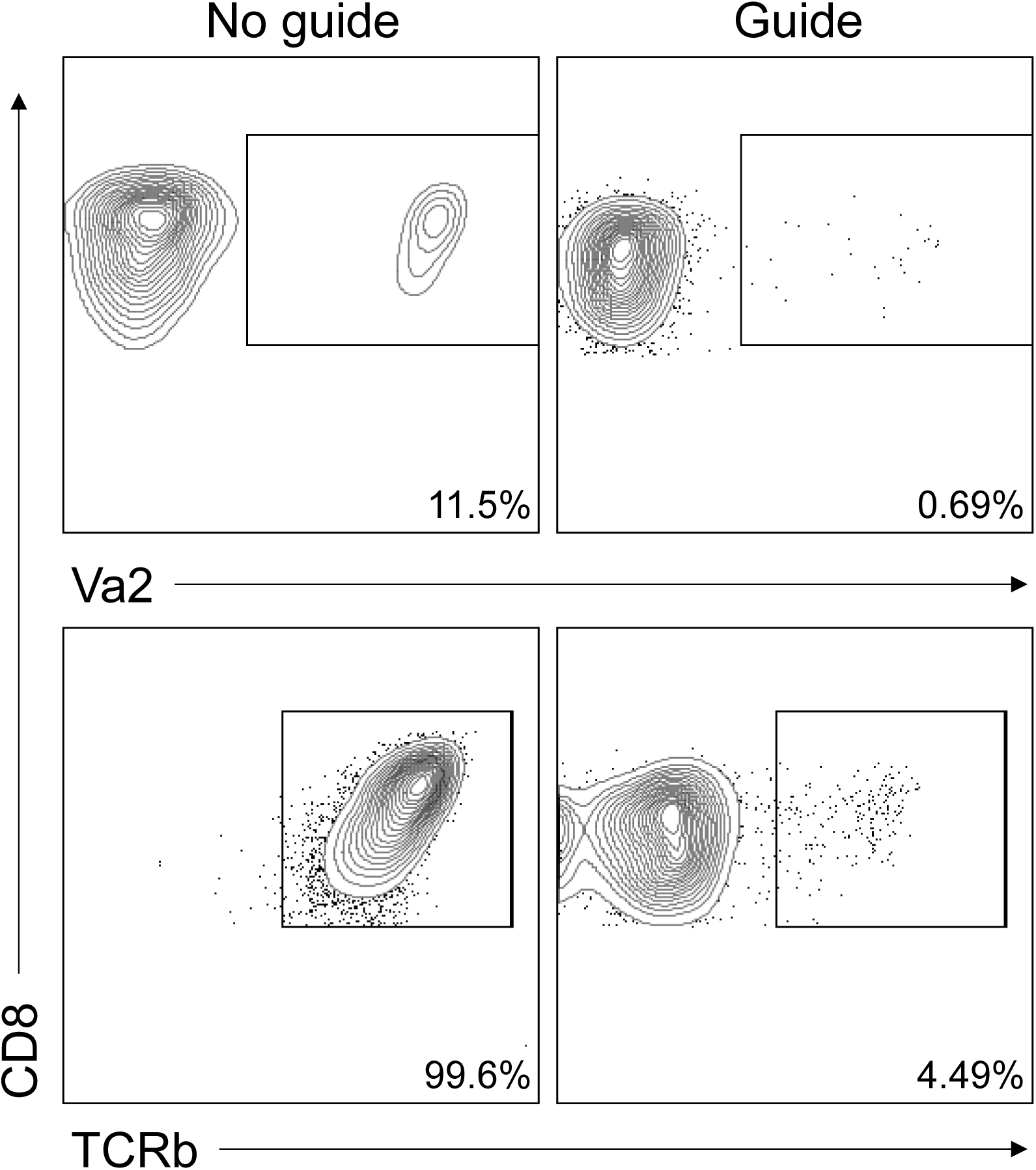
Validation of TRAC and TRBC guide-mediated TCR knockout. (a) Representative flow cytometry plots showing surface expression of Vα2 (on top) and TCRβ (on bottom) following electroporation of primary OT-1 CD8 T cells without sgRNA (No guide) or with a TRAC/TRBC-targeting sgRNA (Guide).

Figure 3 illustrates why efficient removal of the endogenous TCR is necessary, using polyclonal CD8 T cells as an example. Cells receiving virus without CRISPR editing retain endogenous TCR chains while expressing the introduced construct. This creates a scenario where the cell may express the endogenous TCR, express the introduced TCR, express both TCRs (dual specificity), or express a chimeric TCR (i.e. endogenous alpha and introduced beta). It should be noted that all of these possibilities, can negatively impact the purity of the desired clonal population. When CRISPR-mediated TCR knockout is combined with retroviral transduction, the overwhelming majority of transduced cells should lose endogenous TCRβ while a subset will express the Thy1.1 reporter and re-express TCRβ in a manner proportional to retroviral integration. Importantly, similar frequencies of Thy1.1-positive cells are typically observed in the presence or absence of guide RNAs, demonstrating that CRISPR-Cas9 editing may not impair retroviral transduction efficiency.

**Figure 3:**
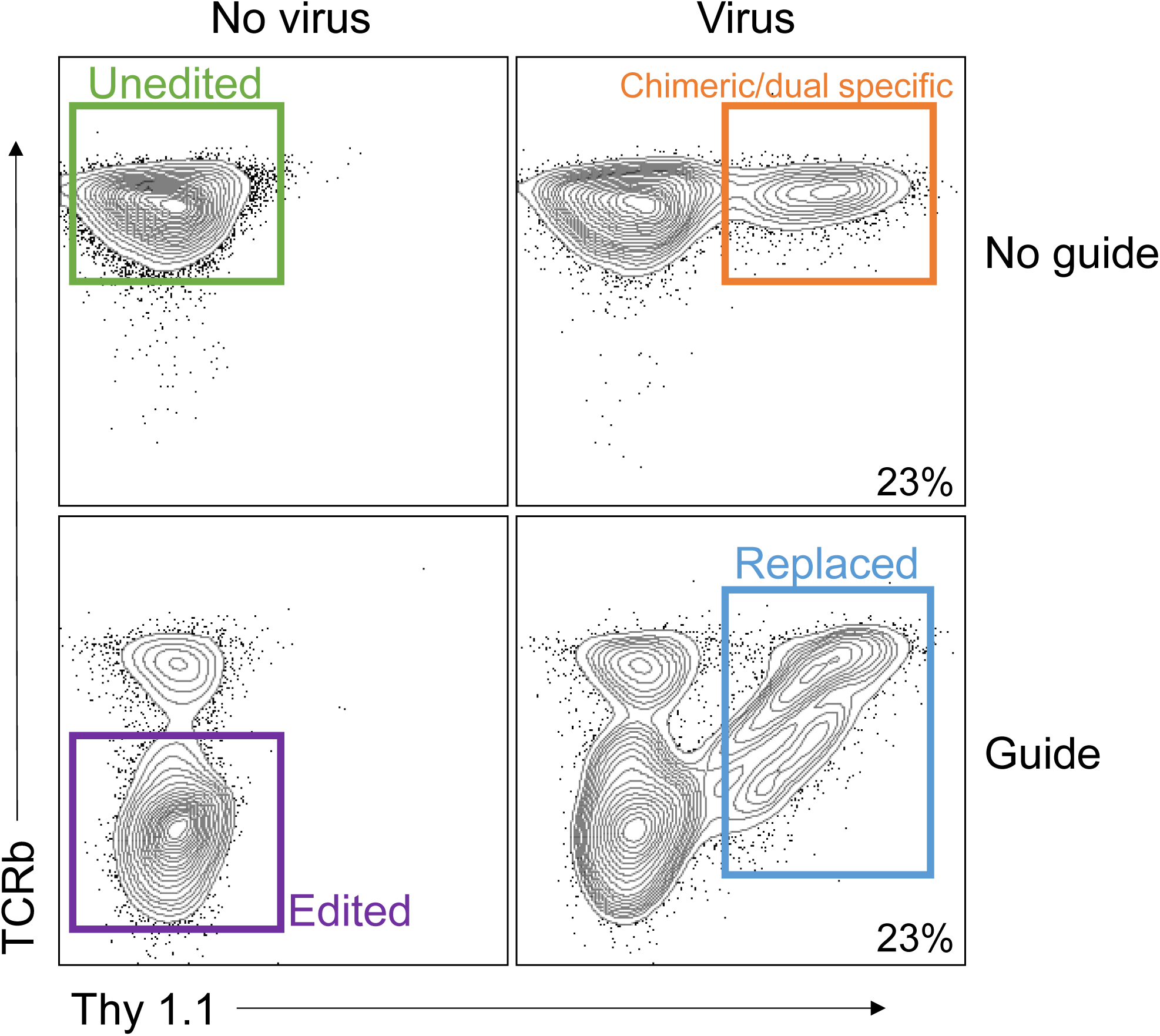
Knockout efficiency of CRISPR-Cas9-mediated disruption of endogenous TCRα and TCRβ expression. (a) Representative flow cytometry plots showing endogenous TCRβ (y-axis) and Thy1.1 reporter expression (x-axis) following CRISPR-Cas9 editing and retroviral transduction of polyclonal CD8 T cells. Cells receiving neither sgRNA nor virus remain unedited (green; TCRβ Thy1.1□). Retroviral transduction in the absence of TCR editing produces T cells with dual specificity and/or chimeric cells co-expressing endogenous TCRβ and the retroviral transduction reporter (orange; TCRβ Thy1.1□). CRISPR editing alone efficiently disrupts endogenous TCRβ expression (purple; TCRβ Thy1.1□). Combining TCR knockout with retroviral transduction generates TCR-replaced cells that lack endogenous TCRβ while expressing the introduced Thy1.1-marked receptor (blue; TCRβ Thy1.1□).

**Figure 4:**
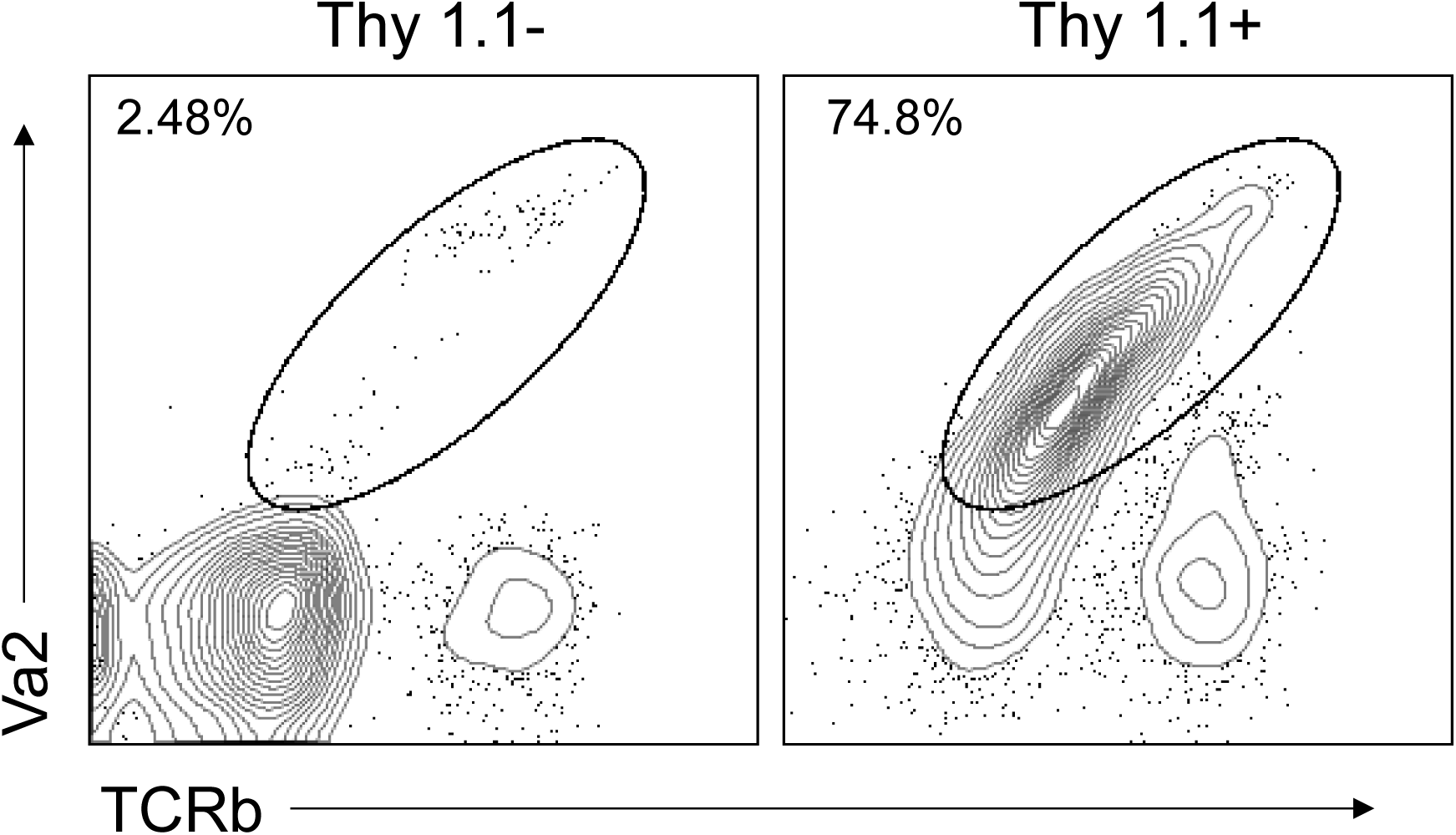
Retroviral transduction efficiency and expression of the introduced P14 TCR following CRISPR-Cas9-mediated TCR editing. (a) Representative flow cytometry plots showing expression of the introduced P14 TCR, identified by Vα2 and TCRβ staining, within Thy1.1 and Thy1.1 populations at day 6. Plots are gated on CD8 T cells and either Thy1.1^+^ or Thy1.1^−^ populations.

Following Thy1.1 positive selection on day 6 post transduction, the majority of Thy1.1+ cells should express the introduced TCR (∼75% or greater). In the case of our p14 TCR, success was identified by proportional co-expression of Vα2 and TCRβ, whereas Thy1.1-cells should contain few P14 TCR-expressing cells. Thus, when the endogenous TCR has been removed, Thy1.1 serves as an effective surrogate marker for enrichment of successfully transduced cells, yielding a highly purified population of engineered TCR-replaced CD8 T cells for downstream applications.

To assess the functionality of engineered P14 CD8 T cells *in vivo*, we edited polyclonal CD45.1+ CD8 T cells, transferred 5 x 10^4^ cells into naïve CD45.2+ B6 recipients, and infected intraperitoneally with 2 x 10^5^ plaque-forming units (pfu) of lymphocytic choriomeningitis virus (LCMV-Armstrong strain). As positive controls, separate animals received 5 x 10^2^ naïve CD8 T cells isolated from P14 TCR transgenic mice (hereafter referred to as Transgenic). Eight days after infection, transferred Thy1.1^+^ splenocytes were analyzed by flow cytometry for expression of markers associated with effector and memory differentiation, including KLRG1, IL-7Rα (CD127), and CX3CR1.

As shown in Figure 5, engineered P14 TCR-replaced and Transgenic cells display identical phenotypes at day 8 post-acute infection. Both populations are dominated by KLRG1^+^ CD127^−^ effector cells, whereas the frequency of CD127^+^ KLRG1^−^ memory precursor cells remains low (Figure 5a-b). Similarly, the majority of cells in both groups acquire a CX3CR1 KLRG1 effector phenotype (Figure 5c-d). The low expression of IL-7Rα in the KLRG1low population is most likely due to the timepoint for analysis, as IL-7Rα increases significantly on KLRG1low CD8 T cells following clearance of LCMV infection (between days 7-10; ^22^). Collectively, these findings demonstrate that CRISPR-Cas9-mediated TCR replacement generates antigen-responsive CD8 T cells that phenotypically resemble conventional P14 transgenic T cells during an acute viral infection. Nonetheless, because the method to generate TCR replaced cells requires activation, the method may impact some downstream functions and phenotypes, and will not be suitable for all applications.

**Figure 5:**
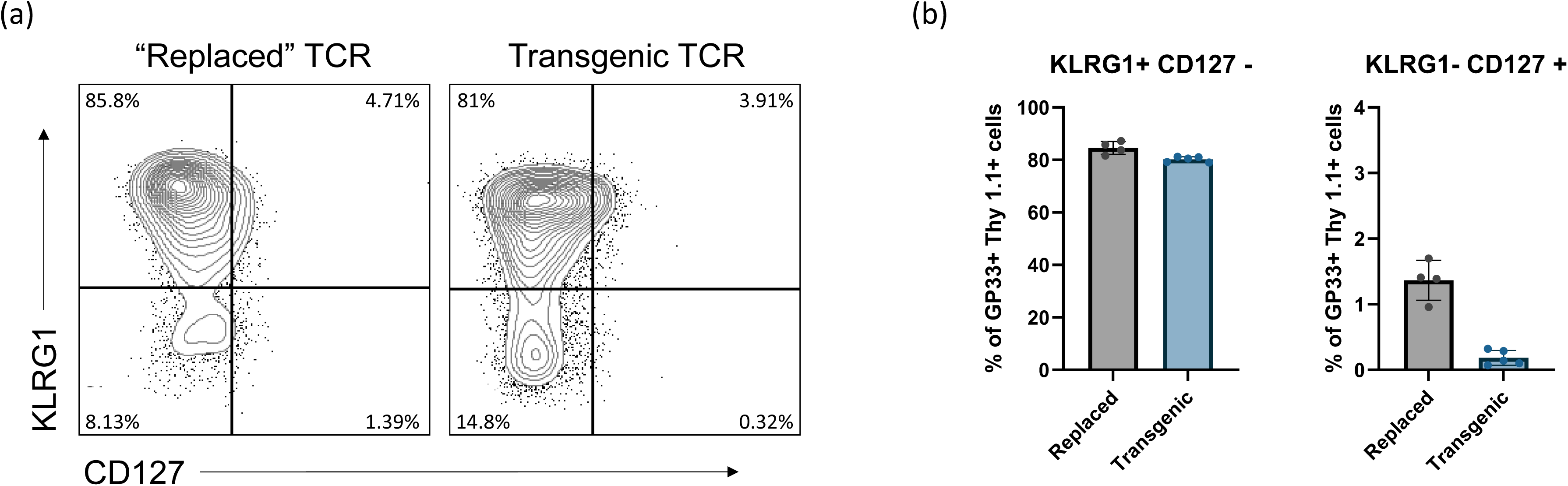
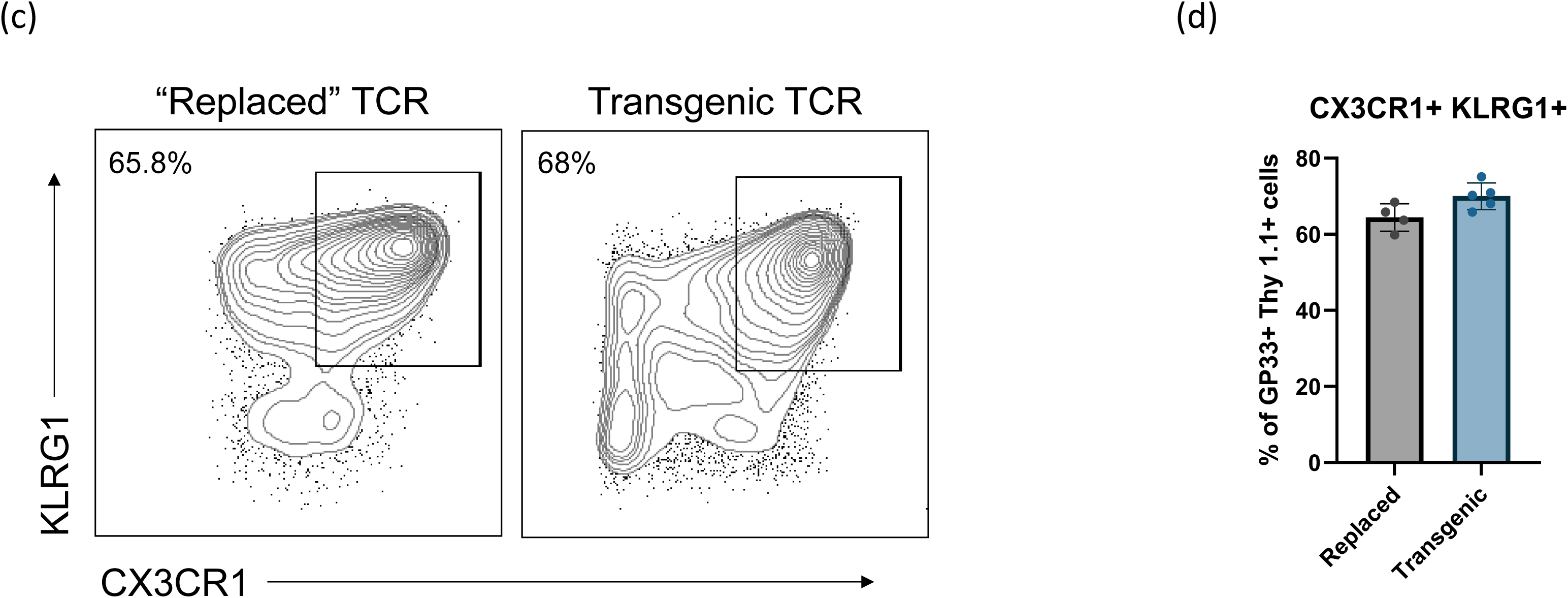
In vivo phenotypic characterization of engineered P14 TCR-replaced and P14 transgenic CD8 T cells following acute LCMV infection. Recipient B6 mice received either engineered P14 TCR-replaced CD8 T cells or CD8 T cells isolated from P14 transgenic mice prior to infection with LCMV. Spleens were harvested and analyzed by flow cytometry 8 days post-infection. (a) Representative flow cytometry plots showing CD127 and KLRG1 expression on transferred P14 TCR-replaced cells and P14 transgenic cells gated on Thy 1.1^+^ GP33^+^ cells. Both populations exhibited a predominantly KLRG1□ CD127□ effector phenotype, whereas the frequency of CD127□ KLRG1□ memory precursor cells remained low (∼1%) in both groups. (b) Quantification of the frequency of KLRG1 CD127 terminal effector cells and CD127□KLRG1□ memory precursor cells among transferred Thy1.1□ GP33□ CD8 T cells from P14 TCR-replaced and P14 transgenic donor populations. (c) Representative flow cytometry plots showing CX3CR1 and KLRG1 expression on transferred P14 TCR-replaced cells and P14 transgenic cells gated on Thy 1.1^+^ GP33^+^ cells. The majority of engineered TCR-replaced cells acquired a CX3CR1□ KLRG1 effector phenotype comparable to that observed in P14 transgenic controls. (d) Quantification of the frequency of CX3CR1 KLRG1 cells among transferred Thy1.1 GP33 CD8 T cells from P14 TCR-replaced and P14 transgenic donor populations.

## Acknowledgements

We thank the Yale Flow Cytometry Core and the Yale Center for Genome Analysis. The graphical summary was created with Biorender.com. We thank the NIH Tetramer Core Facility (contract number 75N93020D00005) for providing (LCMV GP var C41M 33-41 KAVYNFATM H2-Db MHCI) tetramers. We thank the entire Joshi lab for thoughtful discussion and feedback related to this protocol.

## Contributions

N.T., J.A., E.F., developed the protocol. N.T. wrote and revised the manuscript and generated figures. J.A. contributed to figure preparation and manuscript writing. K.C. and N.J. conceived and supervised the work. All authors discussed the results and commented on the paper.

## Ethics declaration

The authors declare no competing interests.

